# An algorithm underlying directional hearing in fish

**DOI:** 10.64898/2025.12.11.693757

**Authors:** Johannes Veith, Ana Svanidze, Benjamin Judkewitz

## Abstract

Humans and other land vertebrates localize sound by comparing the signals in each ear. Even though these differences are virtually absent underwater, fish are still able to sense the direction of sound. In 1975, Arie Schuijf proposed that this ability could arise from a comparison of the particle motion phase and the pressure phase of sound, a prediction which was recently confirmed experimentally for near-field sounds. In natural environments, however, sounds arrive from variable distances, altering the motion-pressure phase relationship. Thus, directional hearing and distance hearing are potentially at conflict. There is currently neither a model nor experimental data for how fish deal with this complexity. Here, we systematically introduce phase differences to the particle motion and pressure components of sound pulses to quantify the directional tuning of startle responses in *Danionella cerebrum*. We find that the fish’s directed startle behavior is both frequency and phase dependent, and introduce a new model that quantitatively predicts the sensorimotor transformation across all observed stimuli. This framework likely extends to other otophysan fishes with evolutionarily conserved hearing apparatus, representing ∼15% of all vertebrate species.

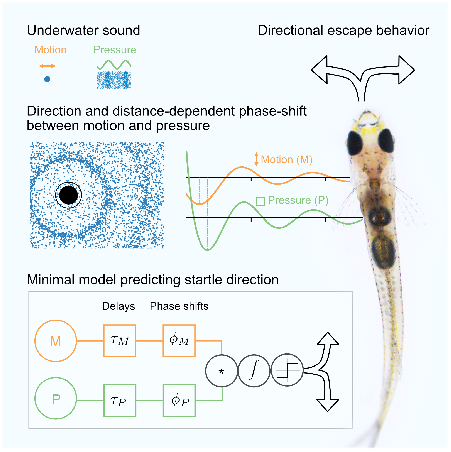

## INTRODUCTION

To survive, animals evolved strategies to escape from sudden sounds. Whether in air or in water, sound is a mechanical wave consisting of pressure and particle motion oscillations (Figure 1A-B). Humans and other terrestrial animals use pressure to infer the direction of sound by sensing it in a spatially extended manner (two eardrums) or by asymmetrically perturbing sound (pinna). While time delays and intensity comparisons between two ears tell sound direction on land, this is unlikely to work for most fish, given the much higher speed of sound and acoustic impedance match to water ^1,2^. Unlike human eardrums which oscillate with only the pressure signal, fish can detect both pressure and particle motion ^3^: In many species, pressure is sensed indirectly, as it sets the compressible swim bladder into motion. This pressure-related motion either re-radiates or is transduced to the inner ear hair cells via the Weberian apparatus ^4–6^. In contrast, particle motion directly stimulates hair cells due to the differential inertia of otoliths and surrounding tissue, rendering it a vectorial sense: When sound emanates from a source, the axis of particle motion aligns with the direction of sound propagation, allowing fish to detect the source’s axis. However, this does not resolve the 180° ambiguity problem ^1^: a sound source on one side of the fish will make hair cells oscillate along the same axis as a source exactly opposite.

**Figure 1.**
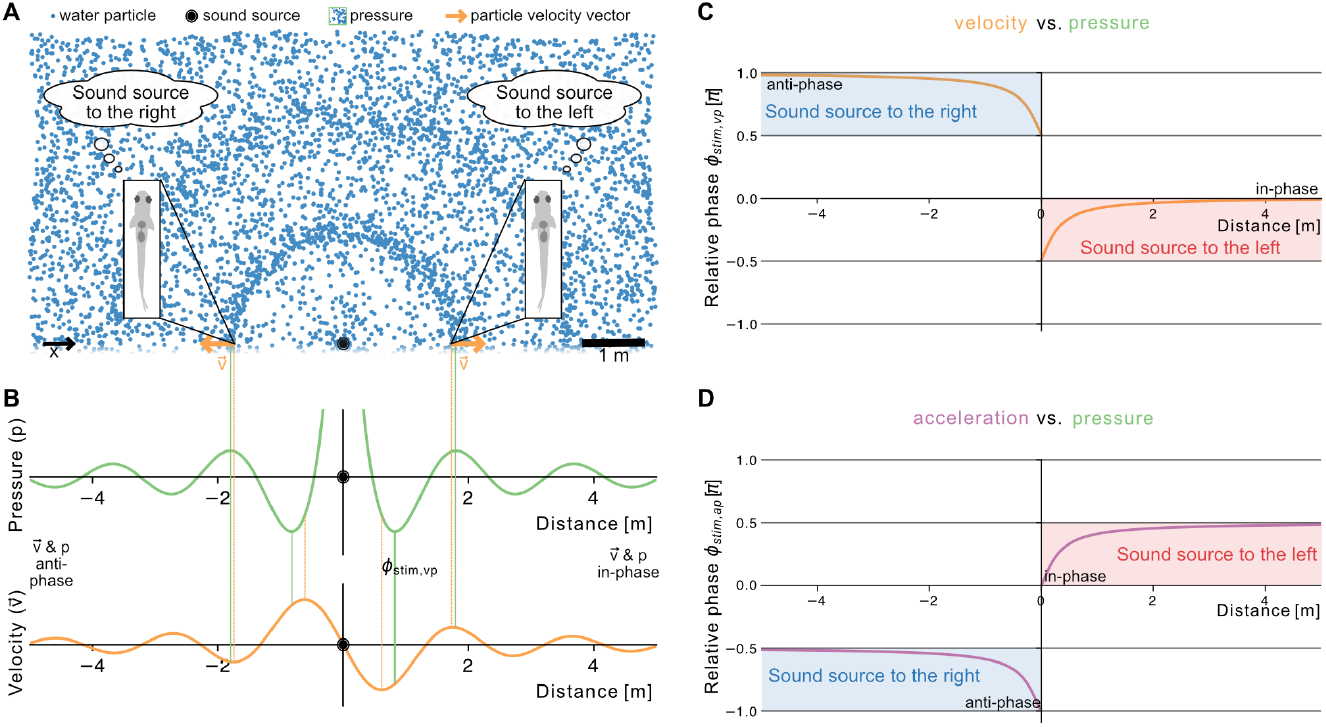
Phase relationship between pressure and particle motion. (A) Schematic illustrating the directional inference challenge in *D. cerebrum*: Neither the pressure nor the particle motion measurements individually resolve the sound source direction. (B) Particle velocity and pressure around a sound monopole. Their relative phase shift Φ_stim,vp_ (between vertical lines), varies with the distance from the sound source. (C-D) The phase-relationship between particle motion and pressure depends on the sound source distance. Nevertheless, it unambiguously falls into two regimes for sound directions. (C) Particle velocity vs. pressure. Far away from the sound source, velocity and pressure are in-phase or anti-phase. (D) Particle acceleration vs. pressure. As particle acceleration is the derivative of particle velocity, it holds that Φ_stim,ap_ = Φ_stim,vp_ + π/2 for sinusoidal motion. Consequently, acceleration is in-phase or anti-phase with pressure very close to the source. (A-D) Shown for a frequency of 800 Hz.

Thus, neither particle motion sensing nor pressure sensing alone explain how small fish can infer the direction of sound. To resolve this dilemma, Arie Schuijf argued in 1975 that fish could combine both sensory channels ^7,8^. He proposed that fish resolve the 180° ambiguity inherent in particle motion by using “a phase analysis” between particle motion and pressure. To make this qualitative statement explicit and to theoretically resolve how the plainfin midshipman fish (*Porichthys notatus*) may be able to swim towards a dipole sound source ^9^, Sisneros and Rogers ^10^ introduced the time-averaged acoustic intensity as a mathematical formulation of Schuijf’s model: 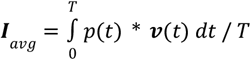. The 180° ambiguity in the oscillatory motion of the vectorial particle velocity **v**(t) is removed by the multiplication with scalar pressure p(t): Far away from a sinusoidal monopole sound source, pressure and particle velocity oscillate in sync. Consequently, the sign of p\***v** defines the direction of the ***I***_*avg*_ vector and captures the direction of sound, effectively performing the phase-analysis proposed by Schuijf. Despite these theoretical efforts, it was long debated whether Schuijf’s model is actually implemented in fish, and the mathematical formulation remained untested.

In our previous work, we confirmed Schuijf’s qualitative prediction and systematically ruled out alternative strategies for directional hearing ^2^. We observed directional escape behavior after acoustically eliciting startles in the miniature transparent fish *Danionella cerebrum* and found that the direction of the startle could be inverted by selectively inverting pressure. Thus, pressure indeed acted as a reference signal to particle motion.

In nature, however, sounds can occur at variable distances. A fundamental property of sound monopoles is that the relative phase between particle motion and pressure is distance-dependent (Figure 1C-D). Therefore, a variable sound source distance poses a challenge to the Schuijf mechanism for directional inference, which is based on a phase-comparison. While the sign of ***I***_*avg*_ still predicts the correct direction across distances, its implementation becomes challenging for near-field sounds (see Supplemental Information): most of the particle velocity signal is phase-shifted by π/2 (Figure 1C) and does not contribute to ***I***_*avg*_, raising questions about the biological plausibility of this mechanism for recent results 2 on directional hearing in the near field.

We concluded that there is currently neither experimental data on the phase-dependency in directional hearing, nor a mathematical model that makes the phase-comparison in Schuijf’s model explicit in a way that explains how direction is extracted despite the fact that phase is distance-dependent.

In this study, we investigate how *D. cerebrum* infers the direction of a sound source across sound source distances and frequencies. To this end, we vary the frequency and phase of parameterized pressure and particle motion waveforms in playback experiments and thereby emulate the relative phases found at variable distances from a sound source. In agreement with the qualitative formulation of Schuijf’s model, we find that the directional inference is explained by this relative phase and not the pressure phase or motion phase alone. By introducing two additional biologically meaningful parameters to a model akin to the time-averaged acoustic intensity – namely phase shifts and time differences between particle motion and pressure – we reproduce the empirical tuning curves of *D. cerebrum* directional startle behavior. The phase and time delays are tuned such that the *D. cerebrum* directional response is optimal for low frequency and nearby sounds, as appropriate for a fast escape behavior.

## RESULTS

### The sound playback paradigm

To understand how the startle direction of *D. cerebrum* is inferred from the relative phase of the particle motion and pressure component of sound, we placed individual fish into an arena (Figure 2A-B) and quantified their lateral (left/right, x-axis) startle behavior in response to a range of acoustic stimuli (Figure S1, S2, Methods). Using multiple speakers and calibrating the impulse response for each speaker we generated precisely controlled combinations of pressure and particle motion targeted to the current fish position (Figure 2C, Methods). This way, we could control both psychophysically relevant components of sound (Figure 2D).

**Figure 2.**
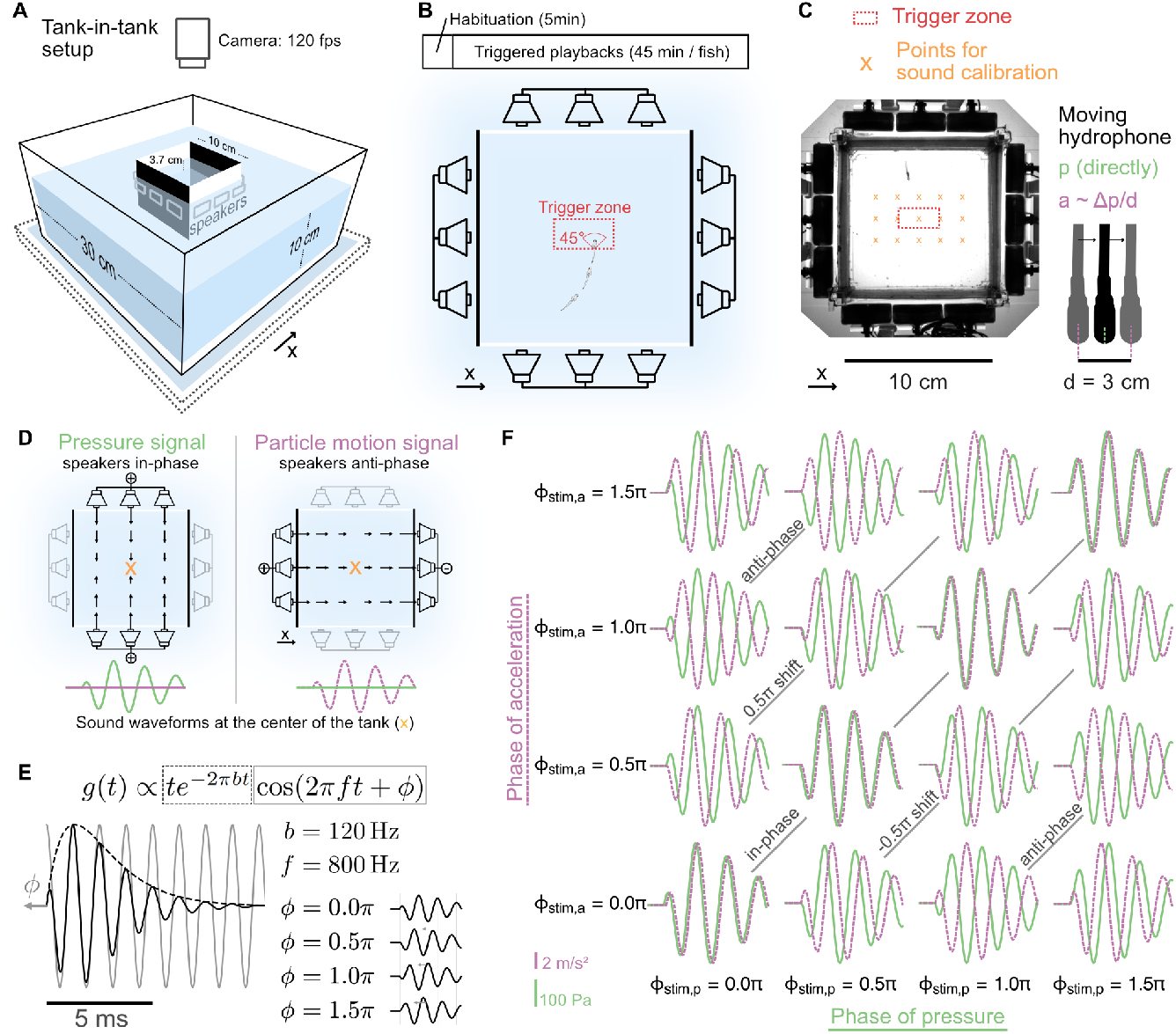
Experimental setup and sound stimuli. (A) Schematic of the behavior setup. Single *D. cerebrum* are placed in the central tank, which is surrounded by underwater loudspeakers. (B) Sound playback is triggered once *D. cerebrum* enters the trigger zone orthogonally to the left-right axis after a minimal timeout interval (> 5s) since the last playback. Two white walls encouraged the fish to swim back-and-forth between them. (C) Positions at which the impulse responses of each set of loudspeakers was measured prior to the experiment. A single hydrophone was moved across the tank automatically to measure pressure. Particle acceleration was calculated from spatial pressure gradients (Methods). (D) Independent control of pressure and particle acceleration. Left: two loudspeakers in-phase create a pure pressure signal at the center. Right: two loudspeakers in anti-phase create a pure particle motion signal at the center. (E) The gammatone filter waveform was used to create pressure and particle acceleration target waveforms at different phases ϕ. The black dashed line describes the envelope, the gray solid line the cosine waveform, and the black solid line the resultant stimulus waveform. (F) The sound stimulus set comprised 16 sounds (experiment 1), spanning a space of pressure and x-acceleration gammatones at four different phases. Peak pressure was 223.87 Pa and peak x-acceleration was 7.54 m/s^2^. See also Figure S1.

In natural environments, the relative amplitude and phase of acceleration and pressure varies with the distance to a sound monopole (Figure 1D). To selectively study the effect of phase shifts on directional hearing, we fixed the relative amplitude between particle acceleration and pressure akin to one at 3 cm and only varied the phases (Methods).

Throughout the study, we utilized parametric sound waveforms that allowed for well-defined phase shifts. For the initial experiment, we designed four different gammatone waveforms at f = 800 Hz with phases 0, π/2, π, and 3π/2 by shifting a cosine under a static envelope (Figure 2E). These four gammatone waveforms were used as templates to define the pressure and x-acceleration waveforms. From their combination we created 16 sound stimuli – mimics of natural motion-pressure phase relationships across different distances and directions (Figure 2F). Based on monopole sound theory, we predicted that the directional bias in startle behavior is a function of the relative acceleration-pressure phase, leading to rightward startles for ϕ_*stim,ap*_ = 0, ϕ_*stim,ap*_ = π/2 and to leftward startles for ϕ_*stim,ap*_ = π, ϕ_*stim,ap*_ = 3π/2. Additionally, we predicted that the phases of acceleration or pressure in isolation have little effect on the directional inference.

### The relative phase determines startle direction

To quantify startle directions across various acoustic stimuli, we separated playback trials into startle and non-startle trials and computed the startle direction as the net x-displacement during the initial 50 ms of the startle trajectories (Figure 3A, Figure S2, Methods). In the first experiment, 2750 playback trials (16 stimuli, 62 fish) yielded 1240 startles, with responses triggered within 17 ms of stimulus onset. We pooled data across fish and calculated the proportion of startles directed rightward for each stimulus. To make our metric for directional bias symmetric, we subtracted 0.5 from the fraction of rightward startles. Thus, directional bias ranges from −0.5 (exclusively leftward startles) to +0.5 (exclusively rightward startles), with 0 indicating no bias.

**Figure 3.**
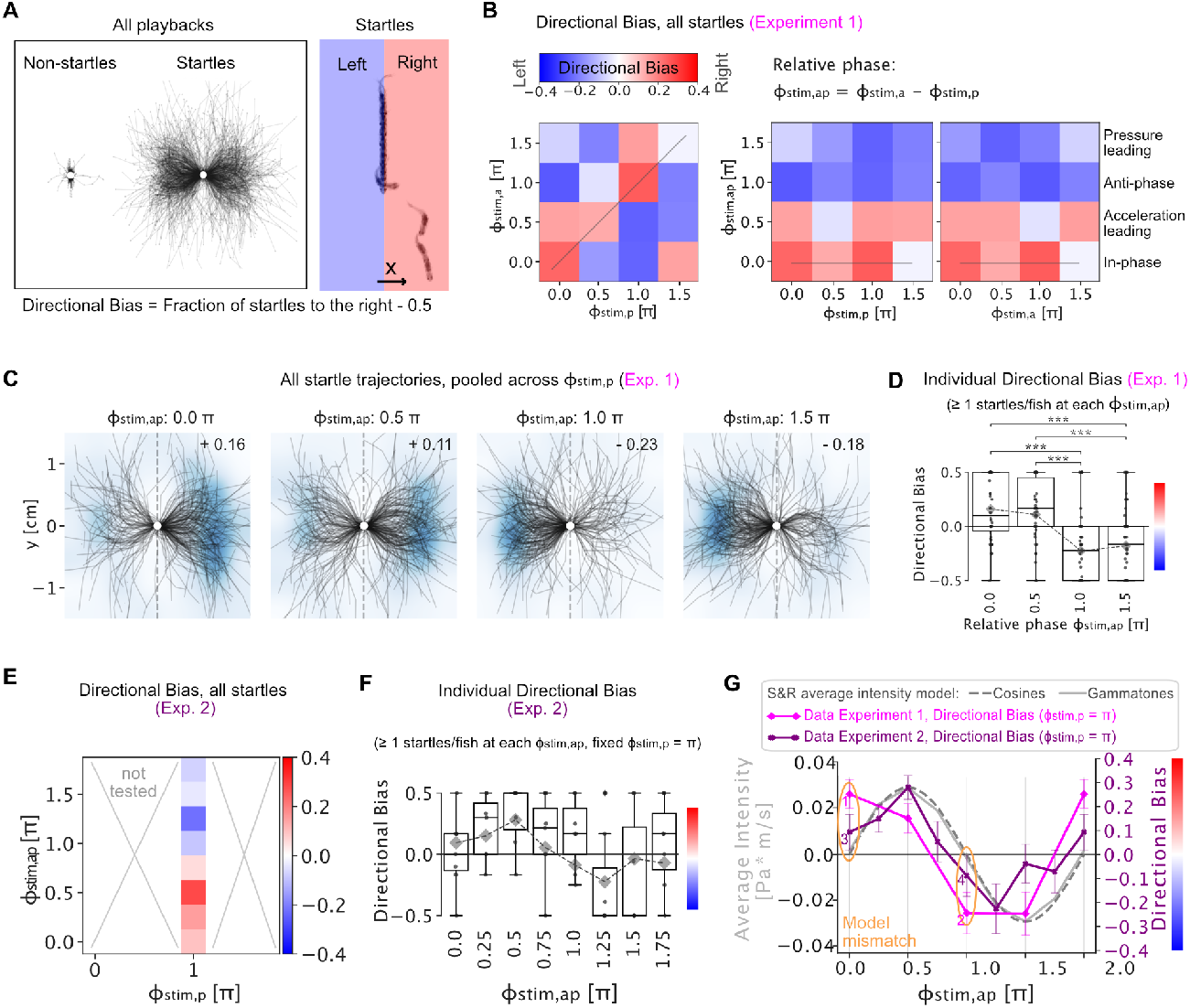
Direction Tuning to Relative Phase. (A) Left: Playback trials fall into non-startle and startle trials, see also Figure S2 and S4B. The centered fish trajectories show the initial 50 ms of the startle response. Right: Directional bias is calculated in the tank’s coordinate system. (B) Heatmaps of directional bias computed across all startles (1240 startles across 62 fish). Left: The invariance across the diagonals reveals that directional bias is a function of the relative phase Φ_stim,ap_. Right: The same data falls into two domains when sorted by Φ_stim,ap_. The thin gray lines highlight identical data in the heatmaps. (C) Centered trajectories for the data shown in (B) pooled across pressure phases as a function of relative phase Φ_stim,ap_. Two-sided binomial tests: 0π: 66% of 267 startles to the right, p = 2 × 10^−7^; 0.5π: 61% of 311 to the right, p = 2 × 10^−4^; 1.0π: 73% of 273 to the left, p = 6 × 10^−14^, 1.5π: 68% of 277 to the left, p = 3 × 10^−9^. (D) Individual directional bias. Box plot: The black dots show directional bias computed for individual fish and the boxes show the median and quartiles. Only fish that had at least one startle in every of the four Φ_stim,ap_ were included (46 of 62 fish). Mean ± SEM: Φ_stim,ap_ = 0.0π: 0.15 ± 0.04; 0.5π: 0.14 ± 0.04; 1.0π: −0.17 ± 0.05; −0.14 ± 0.05, Comparison with paired t-test (Bonferroni corrected with n = 6). ***: p < 5 × 10^−4^. Gray diamonds: average bias across all startles from all fish to sounds with the same Φ_stim,ap_. (E) Directional bias heatmap of experiment 2, probing 8 x-acceleration phases at fixed pressure phase Φ_stim,p_ = 1π, eliciting 701 startles across 90 fish. (F) Individual directional bias. Box plots: Directional bias of individual fish (black dots) for a subset of 8 fish that had at least one startle in response to every sound. Boxes show the median and quartiles. Gray diamonds: average bias across startle from all fish (19 to 71 startles per Φ_stim,ap_). (G) The tuning curves for directional bias computed as average across all startles from experiment 1 and 2 (violet) are overlaid over the analytical model prediction of the Sisneros & Rogers (S&R) model, evaluated for cosine waveforms (gray, dashed line) and for gammatones, i.e. enveloped cosines (gray, solid line). The model captures the general shape of the tuning, but predicts absence of directional hearing at relative phases Φ_stim,ap,_ where directional bias is significant (1: p = 2 x 10-7, 2: p = 6 × 10^−14^, 3: p = 0.3., 4: p = 0.3). The error bars indicate the SEM for a binary variable (computed over 19 to 71 startles).

In line with previous findings ^2^, the directional bias could not be explained by pressure or acceleration phase alone (Figure S3A-B), nor by the ratio of left vs. right speaker amplitude (Figure S3C). Instead, directional bias was determined by the relative phase ϕ_*stim,ap*_ between particle acceleration and pressure (Figure 3B). A pooling across stimuli with identical relative phases ϕ_*stim,ap*_ highlights this strong directional tuning across all startles (Figure 3C) and in individual fish (Figure 3D).

### Directional inference is cosine-tuned

To further resolve the directional tuning to relative phase, we performed a second experiment where we probed the relative phase ϕ_*stim,ap*_ at finer increments by varying the particle acceleration waveforms at 8 different phases, while maintaining a fixed pressure phase (1.0π) (Figure 3E). We elicited 1150 startles across 1638 playbacks in 90 fish. The results reproduced the general trend of directional tuning to relative phase ϕ_*stim,ap*_ and revealed a directional tuning curve reminiscent of a cosine function (Figure 3F).

We next asked whether we could derive this directional tuning curve mathematically. As a starting point we used a model developed by Sisneros and Rogers for the phonotaxis behavior of midshipman fish ^10^. This model posits that fish use the time-averaged acoustic intensity, 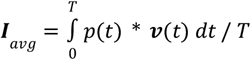, to determine the direction of a sound source. For cosine sounds of angular frequency ω, and particle velocity waveforms that are phase-shifted by ϕ_*stim,ap*_, the expression becomes 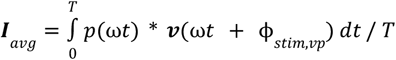. This integral simplifies to *I*_*avg*_ ∝ *cos*(ϕ_*stim,ap*_) (Supplemental Information). Therefore, the time-averaged acoustic intensity follows a cosine with the relative phase ϕ_*stim,vp*_.

In our experiments, we used cosine waveforms under an envelope, and varied the relative phase between particle acceleration and pressure ϕ_*stim,vp*_, and not ϕ_*stim,vp*_. Following from the above analytical derivation, ignoring the envelope and using the fact that velocity is the integral of acceleration (hence ϕ_*stim,vp*_ = ϕ_*stim,vp*_ − π/2), we arrive at the model prediction that startle direction should be a function of *I_avg_* ∝ *sin*(ϕ_*stim,vp*_) (Figure 3G, gray dashed line). A numerical integral of the actual stimulus waveforms – including the gammatone envelope – produces a very similar model curve (Figure 3G, gray solid line).

Thus, a model that was developed to explain a different directional hearing behavior in a different species predicts the cosine-like directional tuning in our startle experiments. Motivated by this realisation, we sought to test whether the time-averaged acoustic intensity *I_avg_* could be turned into a minimal model for directional bias (D) in startle behavior by setting *D* ∝ *I*_*avg*_ (i.e. approximate linearity in the physiological range of sound amplitudes). However, the measured tuning curve appeared systematically shifted against the model *I*_*avg*_, so that the model predicts absence of directional bias, where we observe significant bias, e.g. at ϕ_*stim,ap*_ = 0, and ϕ_*stim,ap*_ = π (Figure 3G, orange ovals). To address this discrepancy, we next introduce a biologically plausible generalization of the model.

### A biologically plausible generalization of the model

Neuronal systems not only detect different sensory signals, but can also implement algorithms to compare them. One such example is the time delay and intensity comparisons between two ears for discriminating the direction of sound in tetrapods ^11^. We therefore consider the option of biophysically and neuronally induced phase shifts and time delays between pressure and particle motion. Along the auditory pathway of *D. cerebrum*, plenty of opportunities exist for creating such comparative signals: Pressure waves set the swim bladder into motion. This motion is transduced via the Weberian apparatus that compresses the perilymphatic sinus, which leads to motion of the sensory macula where hair cells sense displacement ^12,5,6,2^. Thus, a time- and phase-shifted version of pressure may be sensed. Similarly, the particle motion component of sound directly induces differential motion between the otoliths and the surrounding tissue which is sensed as hair cell deflection. This interaction has been modeled to introduce phase shifts ^13,14^. Furthermore, in the neuronal afferent pathway, neurons may preferentially respond to pressure or particle motion, but could introduce additional phase shifts by performing integration or detecting gradients. Additionally, synapses and axons could introduce further differential time delays. To account for these biological effects, we extend the time-averaged acoustic intensity model by introducing a relative phase shift ϕ_*bio,vp*_ and a relative time shift τ_*bio*_ between particle velocity and pressure (Figure 4A).

**Figure 4.**
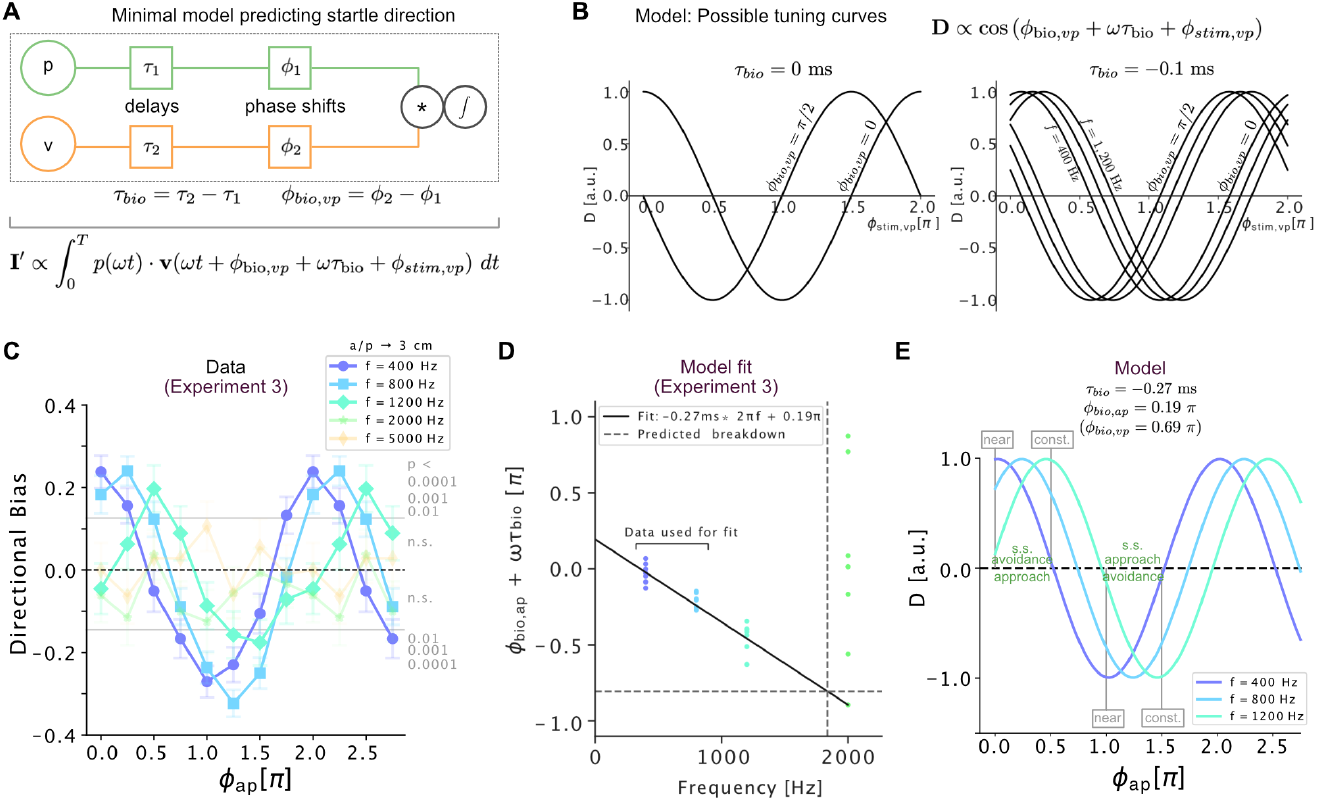
A Mathematical Model for Directional Hearing Behavior. (A) The directional hearing model with biologically acquired delays τ and phase shifts ϕ. The generalized model I’ shifts the particle velocity (v) and pressure (p) inputs by a time difference *τ*_*bio*_ and a phase difference ϕ_*bio,vp*_. Angular frequency ω = 2π*f*. (B) Treated as model for directional bias, *D* ∝ *I*’, D evaluates to cosine tuning for cosine pressure and particle velocity waveforms. Left: Increasing ϕ_*bio,vp*_ shifts the tuning curve to the left. If the time delay τ_*bio*_ is zero, the tuning is independent of the sound frequency. Right: Frequency-dependent rightward shifts emerge from a negative time delay (400 Hz, 800 Hz, 1200 Hz shown). (C) The tuning curves for directional bias from experiment 3 reveal a shift with frequency as predicted from a non-zero time-delay. In total, 3484 startles were evoked across 145 fish. Each data point is computed from 85 ± 30 startles and at least 40 startles. The probability that chance could produce these individual directional biases are reported in grey (p-values for two-sided binomial test with P=0.5). The p-values for individual data points are shown in Figure S5A. Error bars indicate the SEM. The amplitude ratio of acceleration and pressure (a/p) corresponds to a sound source at 3 cm distance. Figure S4 quantifies the startle probabilities (D) We extract the phase of the tuning curves at 400 Hz and 800 Hz and subtract the stimulus phase ϕ_*stim,ap*_. Under our model, ϕ_*bio,vp*_ + ωτ_*bio*_ is the residual phase that is fitted using linear regression. We obtain a negative time delay τ_*bio*_ = − 0. 27 *ms* ± 0. 02 *ms*, and a positive phase shift ϕ_*bio,ap*_ = 0. 19 π ± 0. 04 π. This line correctly predicts the phase of the directional tuning curve at 1200 Hz. At a frequency *abs*(ωτ_*bio*_) > π, leading or trailing become indistinguishable. In line with the experimental data, this breakdown happens at 1837 Hz ± 25 Hz. See also Figure S5C. (E) The fitted model for directional bias *D* ∝ *cos*(ϕ_*bio,ap*_ + ωτ_*bio*_ + ϕ_*stim,ap*_), resembling the data shown in panel C. The directional bias next to a sound monopole can be understood as the addition of a constant term (const., at ϕ_*stim,ap*_ = π/2) and a term that becomes large near a sound source (near, at ϕ_*stim,ap*_ = 0). Figures S5 and S6 show the model fit to the data explicitly. S.s.: sound source.

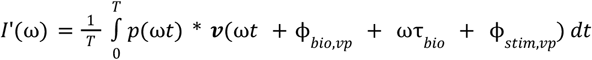

The parameters ϕ_*bio,vp*_ and τ_*bio*_ define the degree to which the particle velocity sense and the pressure sense are phase shifted or delayed with respect to each other prior to their comparison (via the product).

For cosine pressure and particle motion waveforms that are phase-shifted by ϕ_*stim,vp*_, the model’s prediction for directional tuning becomes:

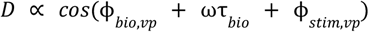

An equivalent formulation of the model in terms of particle acceleration, *D* ∝ *cos*(ϕ_*bio,ap*_ + ωτ_*bio*_ + ϕ_*stim,ap*_), holds true if we define ϕ_*bio,ap*_ = ϕ_*bio,ap*_ + π/2. Under this model, directional bias is maximized if ϕ_*bio,ap*_ + ωτ_*bio*_ = − ϕ_*stim,ap*_ and the fish’s anatomy could theoretically tune these parameters to adapt its behavior to phase shifts ϕ_*stim,ap*_ of natural sounds at specific distances. The model also predicts that for non-zero delays τ_*bio*_, the directional tuning curves should vary with sound frequency (Figure 4B) possibly at the cost of inferring direction correctly across all frequencies.

### The refined model predicts directional tuning

In the previous experiments, we used gammatones at a center frequency of 800 Hz and phase-shifted the x-acceleration waveform against the pressure waveform to obtain a directional hearing tuning curve with relative phase. In our theoretical model we now postulated that there may be both a time delay τ_*bio*_ and a phase delay ϕ_*bio,ap*_ introduced by the fish prior to the comparison of these signals, giving the model more flexibility to fit the data. As time- and phase-delays are indistinguishable at a single frequency, we performed a third experiment using gammatones at multiple frequencies (400 Hz; 800 Hz; 1,200 Hz; 2,000 Hz; 5,000 Hz) while keeping the envelope fixed. We probed 8 different acceleration phases at a single pressure phase (ϕ_*stim,p*_ = π), as before. At the highest frequencies we observed a strongly reduced startle probability (Figure S4A), so we increased the number of playbacks to obtain a similar number of startles for each sound condition (Figure S5A). In this experiment, we obtained 3484 startles from 19590 playbacks across 145 fish. Cosine-like tuning emerged at 400 Hz, 800 Hz, and 1200 Hz, but not at higher frequencies (Figure 4C). Importantly, the *D. cerebrum* directional bias tuning curve was shifted as a function of sound frequency as predicted from a non-zero τ_*bio*_ (Figure S5C). To fit the model, we turned the measured tuning curves into an analytical signal using the Hilbert transform and derived the phase-shift of the tuning curves with respect to ϕ_*stim,ap*_ = 0. Based on the tuning curves at 400 Hz and 800 Hz, we fitted ϕ_*bio,ap*_ + ωτ_*bio*_ and obtained a negative time delay τ_*bio*_ = − 0. 27 *ms* ± 0. 02 *ms*, and a positive phase shift ϕ_*bio,ap*_ = 0. 19 π ± 0. 04 π (equivalent to ϕ_*bio,ap*_ = 0. 69 π ± 0. 04 π). These parameters successfully predicted the 1,200 Hz tuning curve shift (Figure 4D-E). Furthermore, the model correctly predicted that directional inference should break down at *f* > 1837 *Hz* ± 25 *Hz* for which ωτ_*bio*_ >± π and leading and trailing phases cannot be distinguished. We repeated the fit for tuning curves based on startle displacement instead of directional bias and arrived at similar values (Figure S6).

This biophysically motivated adaptation of the Sisneros & Rogers model accurately predicts the directionality of *D. cerebrum* startle responses across frequencies and as a function of relative phase shifts between particle motion and pressure. The model therefore captures a computational algorithm underlying fish directional hearing and can be used to predict which directional valence sounds carry in *D. cerebrum*.

### Directional behavior depends on frequency and distance

These findings have an ecological interpretation. In natural settings, the relative phase ϕ_*ap*_ depends on the sound source distance. Nonetheless, sounds originating from the left and the right fall into distinct ϕ_*stim,ap*_ regimes: Sounds from left create relative phases in the range ϕ_*stim,ap*_ = [0π, 0. 5π] and sounds from right ϕ_*stim,ap*_ = [1π, 1. 5π] (Figure 1D). Across all tested frequencies showing a non-zero directional bias (400 – 1200 Hz), we consistently found that *D. cerebrum* performs startles away from the sound source (Figure 3D, 3F, 4C). Notably, the peak directional bias additionally appeared to be tuned to a specific relative stimulus phase ϕ_*stim,ap, peak*_ (0.03 π at 400 Hz, 0.24 π at 800 Hz and 0.46 π at 1200 Hz) (Figure 4C). Monopole sound theory relates these peak positions to sound source distances (*r* = tan (ϕ_*stim,ap,peak*_) /*k*, wavenumber *k* = 2π*f*/*c*, see Supplemental Information). We find that directional bias is maximal for 400 Hz sounds at phases occurring at distance *r* = ∼6 *cm*, for 800 Hz sounds at *r* = ∼30 *cm*, and for 1200 Hz sounds at *r* = ∼1. 6 *m*. Treating the startle response an escape reaction away from a sound source (*r* = 0), we find that directional bias for nearby sounds is maximized at *f* = 358 *Hz* ± 85 *Hz* and diminishes at *f* = 1277 *Hz* ± 302 *Hz* (Figure S7). Within this framework, the role of a non-zero time shift τ_*bio*_ can be understood as a filter to startle away from nearby sounds with frequencies between 0 Hz and 1277 *Hz* ± 302 *Hz* and most reliably from sounds of frequency *f* = 358 *Hz* ± 85 *Hz*.

To test whether the amplitude ratio of particle motion and pressure – the other distance-dependent quantity (Figure S8A) – could affect these conclusions, we performed a fourth experiment, eliciting 3325 startles in 8885 playbacks across 133 fish. As expected, directional bias decreased with reduced particle acceleration characteristic of more distant sounds. The tuning to the relative phase, however, was unaffected by the amplitude ratio (Figure S8B).

We conclude that the relative phase ϕ_*stim,ap*_ specifically introduces a frequency- and distance-dependent directional bias into the startle trajectory as described by our model.

## DISCUSSION

In this study, we develop a quantitatively validated and generalizable mathematical description of fish directional hearing behavior. We confirm Schuijf’s hypothesis that the relative phase between pressure and particle motion tells the direction of sound and address the complication that the relative phase not only depends on direction, but also on distance. To fit our experimental data, we extend the Sisneros and Rogers model ^10^ by incorporating a time delay τ_*bio*_ and phase shift ϕ_*stim,vp*_ between particle velocity and pressure as tunable parameters. These parameters capture the collective effects of delays and phase shifts between the direct motion sensing pathway and the indirect pressure sensing pathway, allow flexibility in the particle motion quantity being sensed (e.g. displacement or acceleration instead of velocity, or their mixture) and capture the effects of neural integrations or gradient detections. Thus generalized, the model accurately describes the tuning curves for directional startles and explains how the frequency-dependence in directional startle behavior occurs naturally from non-zero time delays. It quantitatively captures strong directional avoidance behavior for nearby, low-frequency sounds (< 1300 Hz, with a maximum at ∼360 Hz), and implies a breakdown of directional discrimination above ∼1800 Hz, matching our experimental observations.

This frequency tuning appears ecologically plausible as predators and abiotic approaching objects tend to emit low-frequency sounds ^15,16^. Two possible explanations for the absence of directional bias at higher frequencies are low-pass filtering in hair cells, apparent in auditory afferents ^17^ and the frequency-dependent reduction in particle displacement (Supplemental Information).

Acoustically-evoked directional startles have been observed in other fish with swim bladder, such as goldfish ^18–24^, herring ^25–27^, angelfish ^28^, cichlid ^21^, and roach ^29^. As *D. cerebrum* shares the Weberian apparatus with ∼66 % of all living freshwater species (comprising ∼15% of all vertebrate species) ^30,31^, we expect our framework to aid quantitative comparative studies across species.

The simple mathematical model reliably transforms sound waveforms into the probability for a directional bias in the startle response, thereby emulating a biological escape circuit. In fish, startles are controlled by Mauthner cells, supported by a broader brainstem escape circuit ^32,33^. A single action potential in the left Mauthner cell triggers a rightward C-bend, catapulting the fish to the right. Inhibitory interneurons amplify even small lateral biases and ensure unilateral activation ^33,34^. Our results suggest that the sound features that drive firing in the escape circuit (measured by startle probability) are independent of the ones that bias the circuit (measured by directional bias): The startle probability is independent of the relative phase between motion and pressure or acceleration phase, but strongly depends on pressure phase and frequency, a finding that is consistent with earlier physiological ^35^ and behavioral reports ^29,36^ (Figure S4, Supplemental Information). Directional bias, in contrast, depends exclusively on the relative motion-pressure phase (Figure 3B). To implement our directional bias model, hair cells could sense pressure and particle motion in a fully or partly unmixed manner ^37^. Auditory afferents could phase-lock their firing on phase-shifted versions of the impeding pressure and particle motion waveforms, introduce time delays, and perform the multiplication in the form of a coincidence detection, thereby unilaterally biasing the circuit. The physiological possibility of coincidence detection is supported by latency measurements in cells of the goldfish brain stem escape network, attesting a minimum onset latency between acceleration- and pressure-evoked postsynaptic potentials below 1 ms ^35^. This latency is compatible with our behaviorally obtained delay estimate of ∼ 0.3 ms, which constrains connectionist models that have been developed for goldfish ^38^. Future studies measuring the activity and phase-locking capacities of auditory afferents and the brainstem escape circuit, complemented by vibrometry measurements and numerical simulations of the biomechanics of pressure and particle motion sensing pathway, could reveal how the mathematical formula is implemented biologically.

In summary, we have found that the integrated product of phase- and time-shifted pressure and particle motion waveforms arriving at the fish’s ear is a reliable predictor of *D. cerebrum*’s directional startle bias. Both the relative phase and the ratio of particle motion and pressure vary with distance, but we identified the relative phase alone to determine the handedness of the startle direction. We observed that *D. cerebrum* startle direction reflects a choice for optimal sound avoidance for nearby and low-frequency sounds. Also in line with the model, *D. cerebrum* does not avoid nearby sounds at higher frequencies as reliably. Given the predictive power and simplicity of the algorithm we described, we expect it to guide and constrain future studies of its biophysical and neuronal implementation.

## RESOURCE AVAILABILITY

### Lead contact

Further information and requests for resources should be directed to and will be fulfilled by the lead contact, Benjamin Judkewitz.

### Data and code availability

Fish trajectory data have been deposited in the G-Node repository at gin.g-node.org/danionella/veith_et_al_2026.

Code used for defining the stimuli, performing the sound calibration, running the experiment, and performing the data analysis is available on GitHub at github.com/danionella/veith_et_al_2026.

## ACKNOWLEDGMENTS

We thank Antonia Groneberg, Kim Kirchberger, Daniil Markov, Maximilian Hoffmann, and Jonathan Bauermann for discussions and for critically reading our manuscript. We thank Arthur N. Popper for constructive feedback regarding the manuscript. We also thank our fish facility team for excellent fish care and experimental support. We acknowledge support by the European Research Council (ERC2021-CoG-101043615), the Einstein Foundation (EPP-2017-413), the German Research Foundation (DFG, projects EXC-2049-390688087 and 432195732) and the Alfried Krupp von Bohlen und Halbach Foundation.

## AUTHOR CONTRIBUTIONS

Conceptualization: JV, BJ; Methodology: JV; Investigation: JV, AS; Visualization: JV; Funding acquisition: BJ; Project administration: BJ; Supervision: BJ; Writing: JV, BJ.

## Declaration of generative AI and AI-assisted technologies in the writing process

During the preparation of this work the authors used ChatGPT-4o in order to improve style and readability. After using this tool/service, the authors reviewed and edited the content as needed and take full responsibility for the content of the published article.

## STAR Methods

### EXPERIMENTAL MODEL DETAILS

#### Fish

All animal experiments and husbandry conformed to Berlin state, German federal and European Union animal welfare regulations. *Danionella cerebrum* were kept at a 14 h / 10 h light / dark cycle in commercial zebrafish aquaria (Tecniplast) with the following water parameters: pH 7.3, conductivity 350 µS/cm, temperature 27 °C. In all four experiments, we used male and female adult fish between 3 and 12 months of age.

## METHOD DETAILS

### Behavioral setup and protocol

In an experiment, a single *D. cerebrum* was swimming inside a tank surrounded by four sets of submerged loudspeakers (4 × 3 Ekulit LSF-27M/SC 8 Ω in a custom water-tight enclosure, Figure 2A). The setup was identical to the one described in more detail before ^2^. For 45 minutes, approximately 6 ms short sounds were triggered when *D. cerebrum* entered the center trigger zone (3 cm x 1.5 cm) with its body aligned orthogonally (within a 45° cone) to the left-right axis and only after a minimal timeout interval (≥ 5s) between playbacks (Figure 2B). The order of acoustic stimuli was randomized for each fish. Two white walls encouraged the fish to swim orthogonally to the left-right axis (there was a 1.6 ± 0.1 times larger absolute displacement along the white-wall axis than the black-wall axis, with standard deviation calculated across the three experiments). In the first experiment, 1240 startles were elicited across 62 fish, in the second, 701 startles across 90 fish, in the third 3484 startles across 145 fish, and in the fourth 3325 startles across 133 fish.

### Sound stimuli

*D. cerebrum* senses sounds as the combination of pressure and particle motion, two tightly coupled components when using a single loudspeaker. Multiple loudspeakers overcome this limitation: In-phase activation of opposite loudspeakers can create a pure pressure stimulus at their center, while out-of-phase activation creates a pure particle motion stimulus (Figure 2D). By balancing in-phase and out-of-phase activations also a desired mix of target pressure and particle motion waveforms can be generated. To increase accuracy and cancel echoes, the impulse responses of each set of speakers was measured at multiple positions in the tank and used to compute loudspeaker activations that could faithfully deliver the target sounds to the fish (Figure 2C, S1). Startles are typically evoked by a sudden loud sound ^18–20,25,26,28,27,21–23,29,24^. As we wanted to investigate the effect of sound phase on directional inference, we reasoned that multiplying a cosine waveform with an envelope that consisted of a steep linear ramp and an exponential decay (Figure 2E) could give us fine control over the phase of a waveform and elicit startles. This waveform is also known as a second-order gammatone filter.

The same envelope was used in all experiments, but the phase and the frequency of the underlying cosine was varied. The target peak amplitudes (pressure: 223.87 Pa, x-acceleration: 7.54 m/s^2^, y-acceleration: 0 m/s^2^) were akin to the amplitude ratio expected next to a sound monopole at 3 cm distance.

Experiment 1 (16 stimuli):

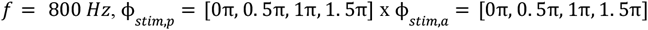

Experiment 2 (8 stimuli):

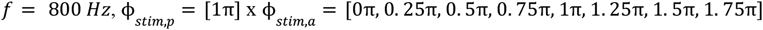

Experiment 3 (40 stimuli):

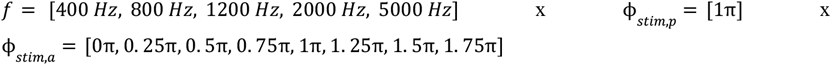

In the fourth experiment, the amplitude ratio of x-acceleration and pressure was varied at a fixed pressure, p = 223.87 Pa.

Experiment 4 (32 stimuli):

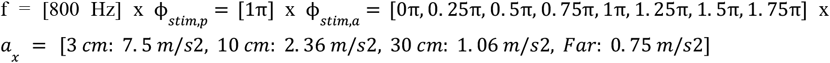

Using the impulse-response based method described in the following paragraph, the targeted sound waveforms in terms of pressure, x-acceleration were faithfully delivered and the y-acceleration was below 2 m/s^2^ (Figure S1). The vertical acceleration (orthogonal to the x- and y-acceleration) was uncontrolled. A triaxial acceleration sensor was used to nevertheless get an estimate (Triaxial ICP, Model 356A45, PCB Piezotronics, acquired with NI-9231 sound and vibration module, National Instruments). Multiplication of the acceleration sensor measurements with a factor 2.4 (to account for the impedance mismatch of sensor and water) reproduced the x- and y-acceleration waveforms that were derived from spatial pressure gradients. The untargeted z-acceleration was measured to be similar to the x-acceleration with peak accelerations up to 8 m/s^2^.

#### Impulse response-based sound targeting

Reverbs can strongly distort the sound waveform created by the loudspeakers. To nevertheless create the same sounds at central positions inside the inner tank, the tank’s reverbal influence was actively cancelled with a method described before ^2^. To this end, the impulse responses for all four speaker triplets were measured prior to the experiment. Using a motor, a hydrophone (Aquarian Scientific AS-1, preamplifier: Aquarian Scientific PA-4, acquisition: NI-9231 sound and vibration module, National Instruments) was moved across a 5 x 5 grid with 1.5 cm spacing in the loudspeaker plane. The acceleration was calculated from the spatial pressure gradient 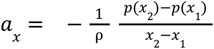. By playing back delta pulses a total of 300 impulse response kernels that described the pressure, x-acceleration and y-acceleration impulse responses (3) for each loudspeaker (4) at all 5 x 5 positions (25) were obtained.

In the following, the sound targeting method is described for one position.

Let *k*_*i,p*_ be the pressure impulse response kernel, 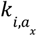 be the x-acceleration impulse response kernel, and 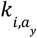 the y-acceleration impulse response kernel for the *i*th speaker. Using *M* speakers with signal *s*_*i*_, pressure and acceleration can be predicted through convolution (*):

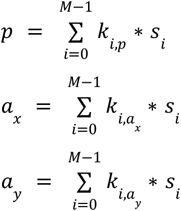

In the Fourier domain, utilizing the convolution theorem, these become a system of equations for each Fourier component *l*.

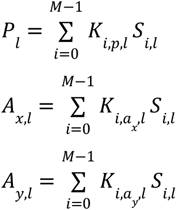

Based on the Fourier components of the target waveforms, *P*_*l*_, *A*_*x,l*_, and *A*_*y,l*_, and the Fourier components of the impulse response kernel *K*_*i,p,l*_,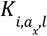, and 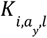, the system of equations can be solved for the Fourier components of the speaker signals *S*_*i,l*_ as long as *M* ≥ 3 and the kernel components are non-zero and non-identical. The time-domain signal for the *i*th speaker is then given by the inverse Fourier transform using components *S*_*i,l*_.

To increase robustness of the solutions (e.g., to avoid speakers canceling themselves unnecessarily and to limit speaker amplitude), speaker signal waveforms were forced to become similar to the target waveform. This was implemented by solving the system of equations with a least square solver (scipy.optimize.lsq_linear) with bounds − *B*_*i,l*_ < *S*_*i,l*_ < *B*_*i,l*_. The bound *B*_*i,l*_ was computed as a rescaling of the absolute Fourier components of the target pressure waveform *P*_*l*_

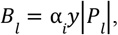

where γ is fixed (scaling pressure to voltage) and α_*i*_ is a rescaling parameter set independently for each speaker to give additional control over active speakers. In these experiments, we set α_1_ = α_2_ = α_3_ = α_4_ = 1. Thus, the bounds set for each speaker were identical.

After conditioning, all computed speaker signals were band-pass filtered between 200 Hz and 1600 Hz (Exp 1 & 2) or 200 Hz and 8000 Hz (Exp 3 & 4) to avoid activating the lateral line. Previous experiments with an identical high-pass filter and similar stimuli had already shown that the capability to exhibit directional behavior after playback is independent of the lateral line function ^2^.

### Behavioral analysis

The fish were tracked using SLEAP ^39^. Some playbacks resulted in a sharp increase in speed (Figure S2A). The speed after playback was averaged over a 25 ms window and revealed a bimodal distribution separating startle trials from non-startle trials (Figure S2B). Based on this, all trials with average speed above 17 cm/s were classified as startle trials (Figure S2C). A heatmap over the fish displacements at multiple delays, showed that the lateral (left/right) displacement was most pronounced 50 ms after startle initiation (Figure S2D). To quantify the directionality of startle trials, we calculated the net displacement along the x-axis within these initial 50 ms. The directional bias was calculated as the fraction of rightward startles minus 0.5. Thus, a directional bias of 0.5 is reported if 100% of startles were to the right and a directional bias of −0.5 if 100% of startles were to the left. Directional bias and directional displacement were highly correlated (Figure S6B). With most fish contributing startles, we computed directional bias by pooling startles across all fish (Figure 3B-C, 3E, 4C).

In some cases (Figure 3D, 3F) – wherever sufficient number of startles per stimulus category and fish could be elicited – directional bias was computed for individual fish after filtering the data for fish that had at least one startle in each stimulus category that was compared. Note, that this per-fish comparison automatically selects for fish with many startles and therefore comes at the cost of introducing an implicit bias to fish that either swam a lot across the trigger zone and/or were likely to startle. Nevertheless, both metrics agreed throughout. Thus, the directional bias that was observable in the mean across startles was also observed at the individual level and did not stem from a few expert fish.

## SUPPLEMENTAL INFORMATION

### Sound monopole theory

The radial pressure component of a sinusoidal monopole sound source is given by ^40^

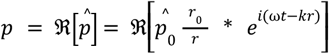

where ω = 2π*f* is the angular frequency, *k* = ω/*c* the angular wavenumber, *p*_0_ the pressure amplitude at radius *r*_0_, and *c* the speed of sound. The hat indicates that the amplitude is complex and ℜ is the real part operator.

The radial pressure gradients will result in radial acceleration, as described by the Euler momentum equation ^40^ with medium density ρ:

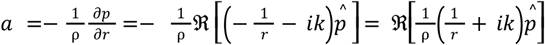

From the particle acceleration, we can also derive particle velocity and particle displacement.

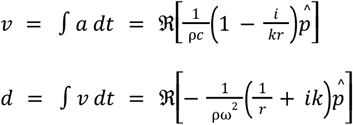

Consequently, the relative phase between radial pressure and radial particle motion are given as:

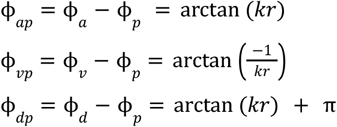

### Predictions of the Sisneros and Rogers model

Sisneros and Rogers propose the time-averaged acoustic intensity as a mathematical formulation of Schuijf’s model ^10^

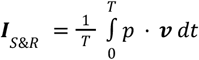

In our experiments, we used pressure and x-acceleration waveforms at multiple phase-shifts. These phase-shifts included those phase shifts that occur radially next to a sound monopole, as derived above. We defined the pressure and x-acceleration waveforms as gammatones:

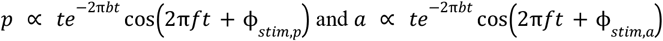

To simplify, we ignored the envelope, *t*e^−2π*bt*^, derived particle velocity from particle acceleration and computed *I*_*S&R*_ over one period *T* = 1/*f*. Thus, we compute the integral for

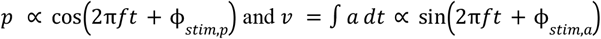

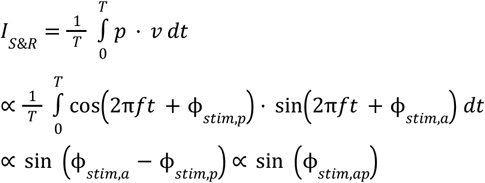

The model predicts a sinusoidal directional tuning with the relative phase between particle acceleration and pressure.

Solving the integral numerically with *T* → ∞, including the gammatone envelope, *t*e−2π*bt*, produces a very similar curve (Figure 3G).

### Generalization of the Siseros and Rogers model

We generalize the Sisneros & Rogers model ^10^ by introducing a relative time shift τ_*bio*_ and a relative phase shift ϕ_*bio, vp*_ between particle velocity and pressure and explicitly apply it to sinusoidal waveforms with angular frequency ω = 2π*f*. This generalization can implicitly capture whether particle displacement, particle velocity or particle acceleration or any integral or derivative of pressure is sensed by the fish. The two introduced parameters allow the fish to tune the directional inference to a specific motion-pressure phase relationship in the stimulus.

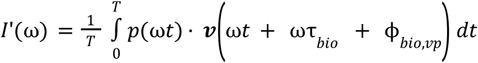

With this modification, we recompute the model’s prediction for the approximated sound stimuli

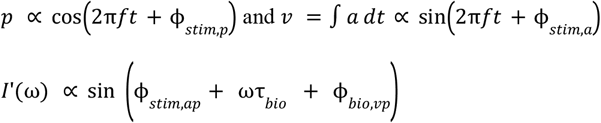

Which, using the definition ϕ_*bio,vp*_ = ϕ_*bio,vp*_ + π/2, equates to

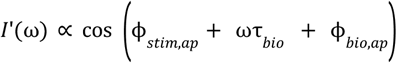

The introduced time delay and phase shift (τ_*bio*_, ϕ_*bio,ap*_) now control how directional inference is tuned to the relative phase between particle acceleration and pressure in the stimulus (Figure 4B). Next, we calculate *I*’ next to sound monopoles.

### Directional inference next to sound monopoles

Next to a monopole, pressure and particle velocity are related as derived above

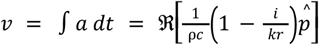

Thus, for sinusoidal waveforms of angular frequency ω, the model evaluates to

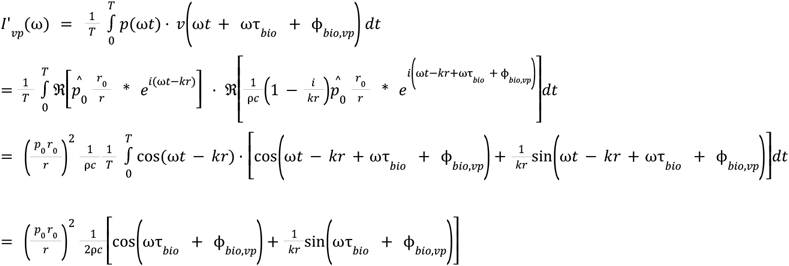

In the near field, as *kr* ≪ 1, *I*’ is dominated by the second summand. The fact that our generalized model equates to the Sisneros & Rogers model if ωτ_*bio*_ + ϕ_*bio,vp*_ = 0, for which that second term vanishes, illustrates that the Sisneros & Rogers model is computing direction only from the far field component. Our generalization allows directional inference based on the dominating near field term, close to a sound source.

For completeness, we also provide the equivalent results for particle acceleration and particle displacement

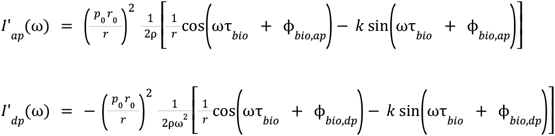

Together with the measured τ_*bio*_ and ϕ_*bio,dp*_ = ϕ_*bio,ap*_ + π, we used *D* ∝ *I*′_*dp*_ to predict the directional inference for different frequencies and across distances (Figure 4F).

### Whether to startle

Here we summarize analyses on the startle probability that – in addition to the algorithm for a directional bias – could inform a neural circuit model. We wondered whether the strong tendency to avoid low frequency and nearby sounds goes along with a higher probability to startle in the first place. If *D. cerebrum* would extract distance information from the relative phase between particle acceleration and pressure and startle more likely to nearby sounds, we would expect the highest startle probability to sound stimuli with Φ_*stim,ap*_ = 0.0π, and Φ_*stim,ap*_ = 1.0π. Instead we find that the startle probability is uniform over the relative phase Φ_*stim,ap*_ (Figure S4C). Startle probability appears much stronger influenced by two other factors. First, there is a steep drop off between sound frequencies of 1200 Hz and 2000 Hz (Figure S4A). Second, startle probability is strongly affected by the pressure phase alone. In line with sporadic reports in other fish ^29,35,36^, *D. cerebrum* appears to be capable of pressure-phase-hearing and startle much more likely to gammatones with pressure phase ϕ_*stim,p*_ = π than to the opposite phase, independent of acceleration phase (Figure S4B).

## SUPPLEMENTAL FIGURES

**Figure S1.**
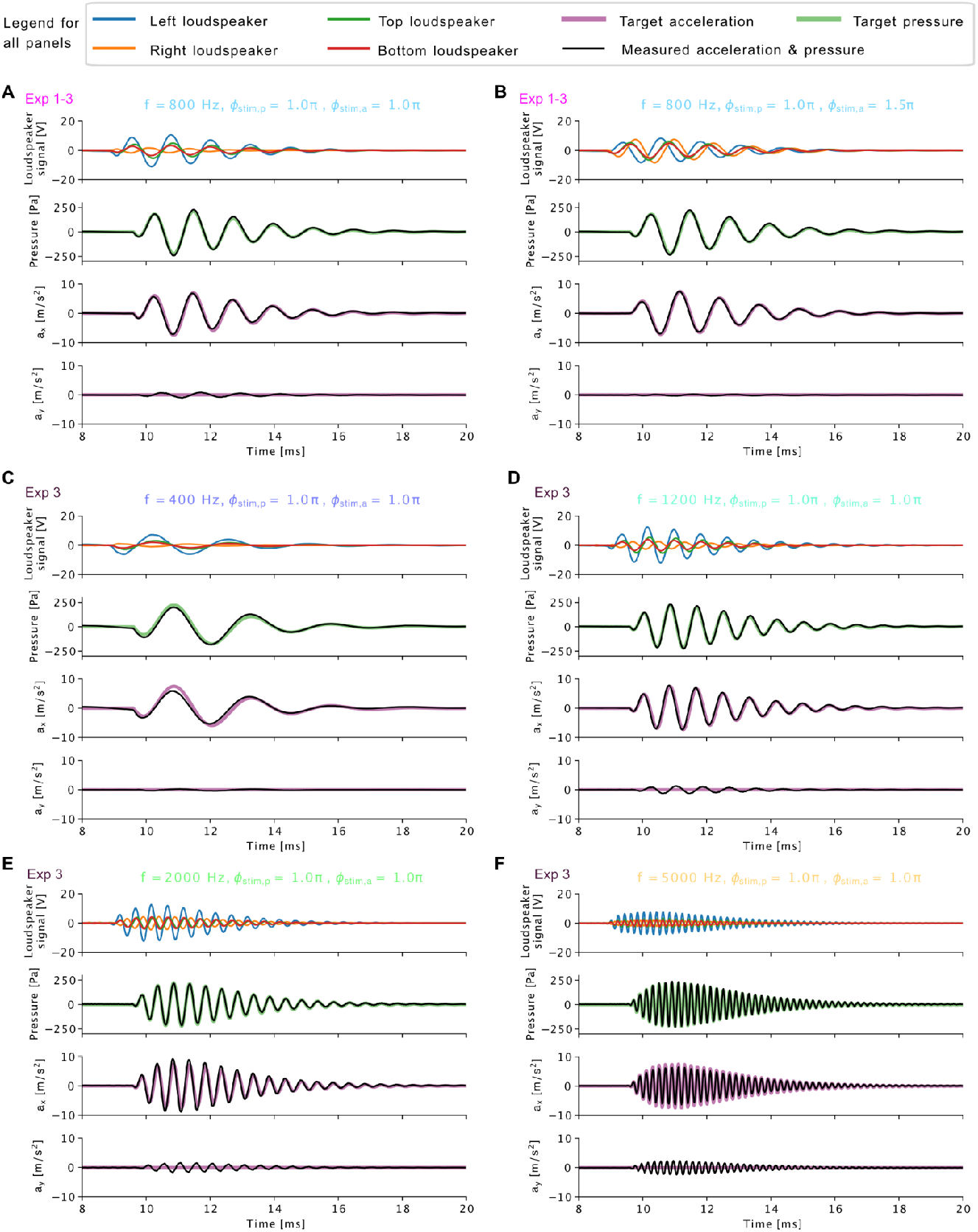
Sound calibration. (A-F) The accuracy of the sound targeting method is shown for six representative sounds. Prior to the experiment, a single hydrophone was moved across the inner tank by a motor to calibrate the sound field (Figure 2C). The impulse responses for each loudspeaker and for each position in the tank were used to calculate loudspeaker signals that could faithfully deliver desired pressure and acceleration waveforms to a target location (Methods). Acceleration (a_x_, a_y_) was calculated from the spatial pressure gradient (Methods). The recorded signals (black lines) matched the target sound waveforms for pressure (green lines) and acceleration (purple lines). Here, we show sound targeting results for the center of the trigger zone. Off-center sound targeting within the trigger zone had comparable accuracy. The constancy of the sound calibration was validated in-between and after the experimental sessions.

**Figure S2.**
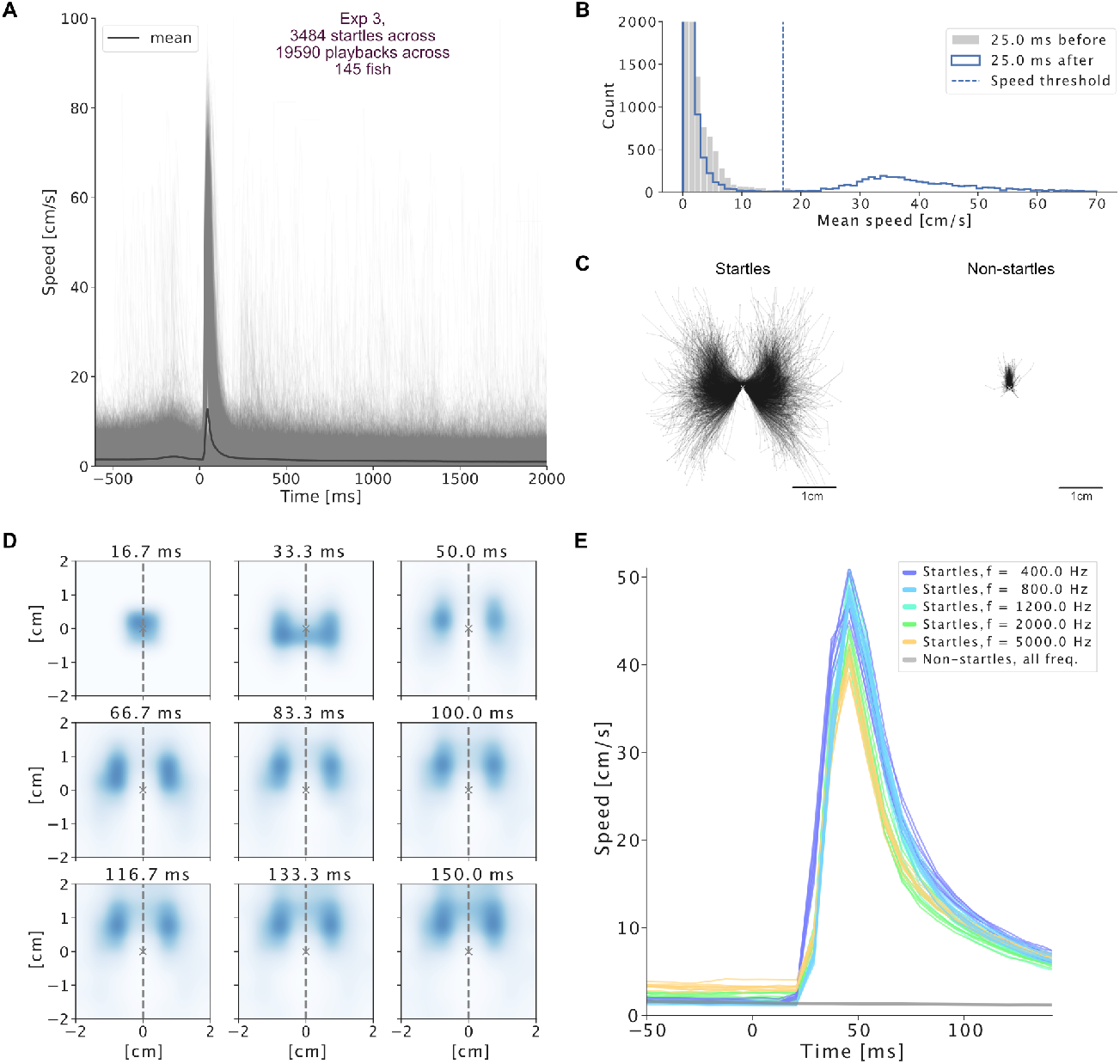
Startle characterization. (A) Fish head speed aligned by sound trigger for each playback. The mean over all playback trials (startles and non-startles) shows a precisely timed increase in speed within 17 ms of sound arrival. (B) After playback, at the time of startle onset, the speed histogram is bimodal at 17 cm/s. Playback trials with an average speed above 17 cm/s were classified as startles, the others as non-startles. (C) Centered trajectories of startles and non-startles. The trajectories are rotated such that the fish faces up at startle initiation. The first 50 ms of the trajectory are shown. (D) The smoothed histogram over endpoint positions after startle initiation at variable time delays reveals that the directional decision is most prominent after 50 ms/s. Afterwards the fish continue swimming toward the front. (E) Time delays and startle speeds vary slightly by frequency.

**Figure S3.**
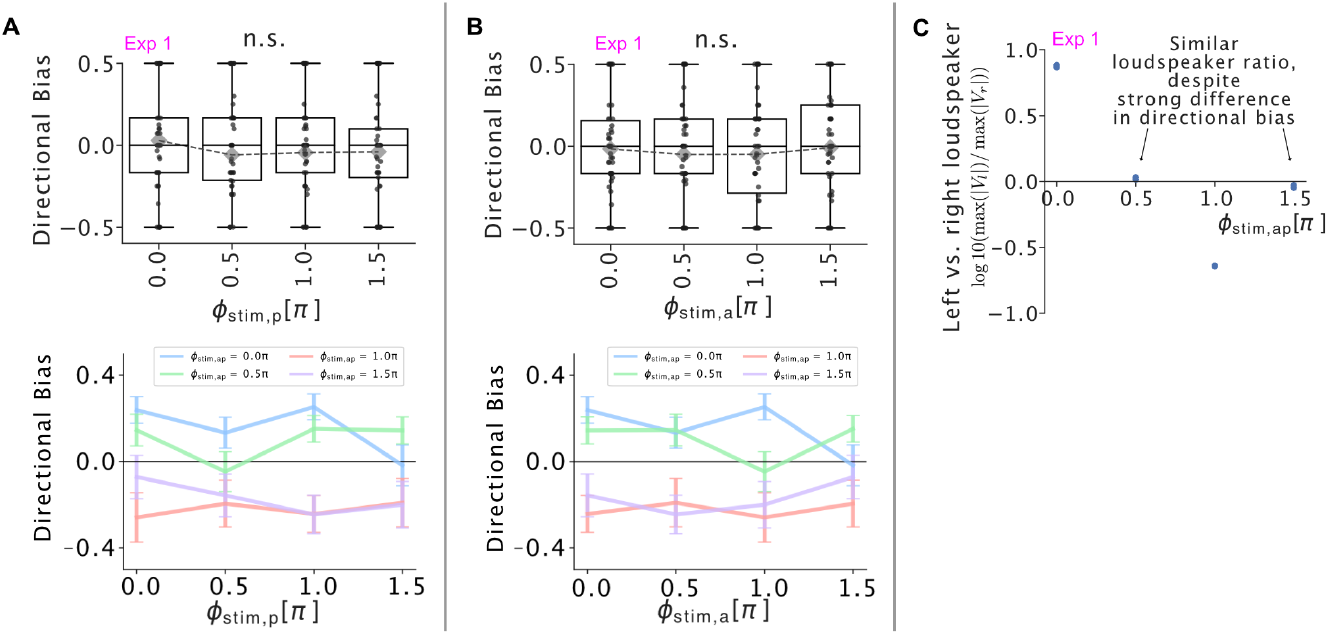
Variables not explaining directional bias. (A) The directional bias is independent of the pressure phase. (B) The directional bias is independent of the acceleration phase. (C) The strong difference in left-right directional bias observed for Φ_*stim,ap*_= 0.5π and Φ_*stim,ap*_= 1.5π in Figure 3D are not explained by differential left vs. right loudspeaker levels. (A-B) Top: Dots show directional bias for individual fish filtered for fish that had at least one startle in response to each pressure phase (A) or acceleration phase (B). Paired t-tests reveal no significant (n.s., p > 0.05) differences. Boxes show the median and quartiles of the individual data points. Diamonds show the average directional bias across all startle trials. Bottom: Error bars indicate the SEM.(A-C) Data from experiment 1 (f = 800 Hz), same as in Figure 3B-D.

**Figure S4.**
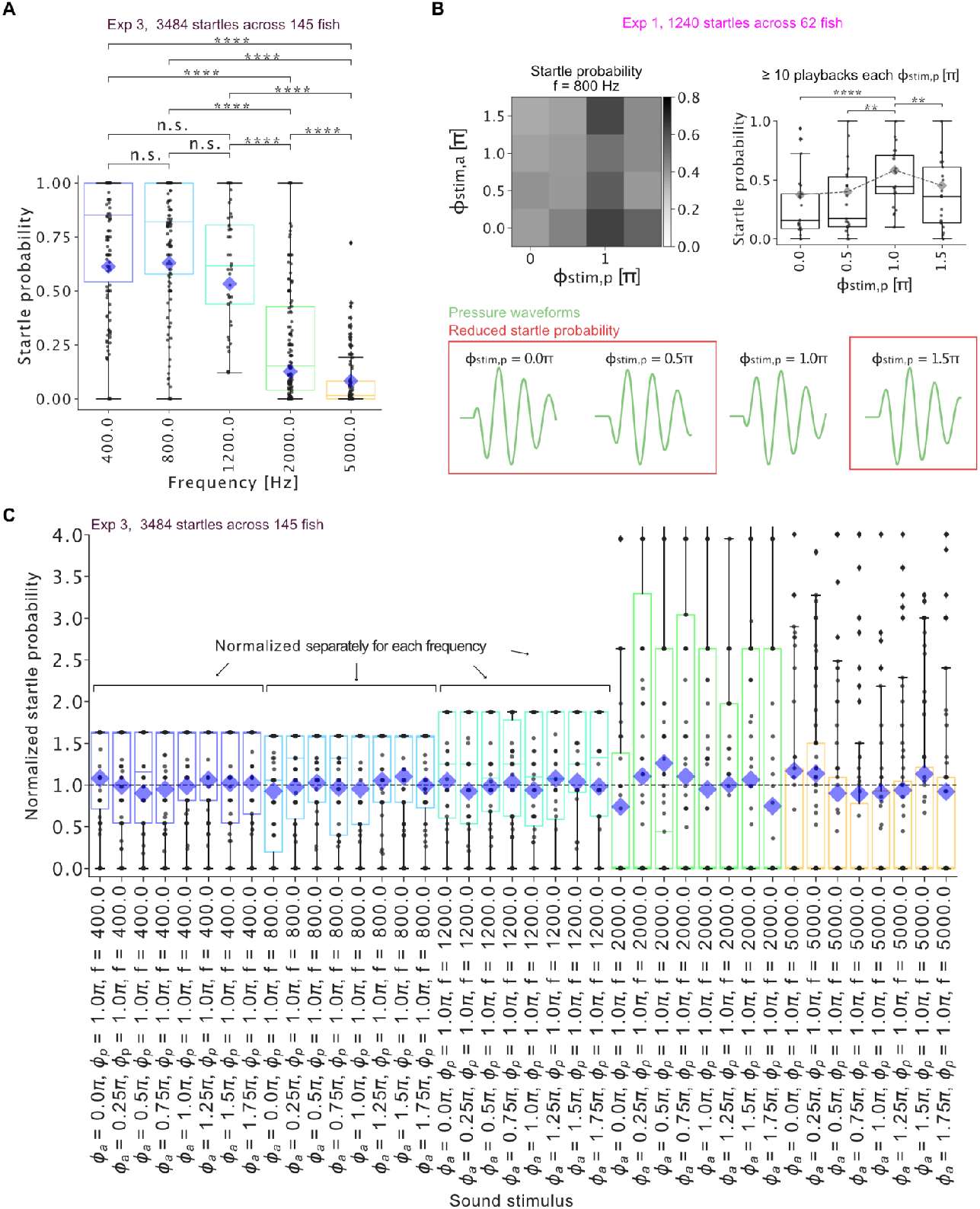
Startle probabilities at different sound conditions. (A) The startle probability drastically decreases with frequency at fixed pressure and acceleration amplitudes. We used an early estimate of these startle probabilities to play back high frequency sounds at a proportionally higher probability to obtain a similar number of startles across sound frequencies. Independent samples t-test, Bonferroni corrected for 10 comparisons, ****: p < 2 × 10^−7^. (B) We observed that a specific pressure waveform (Φ_stim,p_= 1π) elicited startles much more likely. This finding is in line with earlier reports on pressure polarity sensitivity in fish hearing systems ^29,35,36^. Paired samples t-test, Bonferroni corrected for 6 comparisons, ****: p < 0.0001, **: p < 0.003, differences between other pairs are not significant. (C) In contrast, the x-acceleration phase (and relative phase between x-acceleration and pressure) did not affect the startle probability, despite carrying information about the sound source distance. (A-C) Black dots show the (normalized) startle probability of individual fish. Box plots indicate the median and quartiles. Diamonds indicate the mean startle probability across all playbacks.

**Figure S5.**
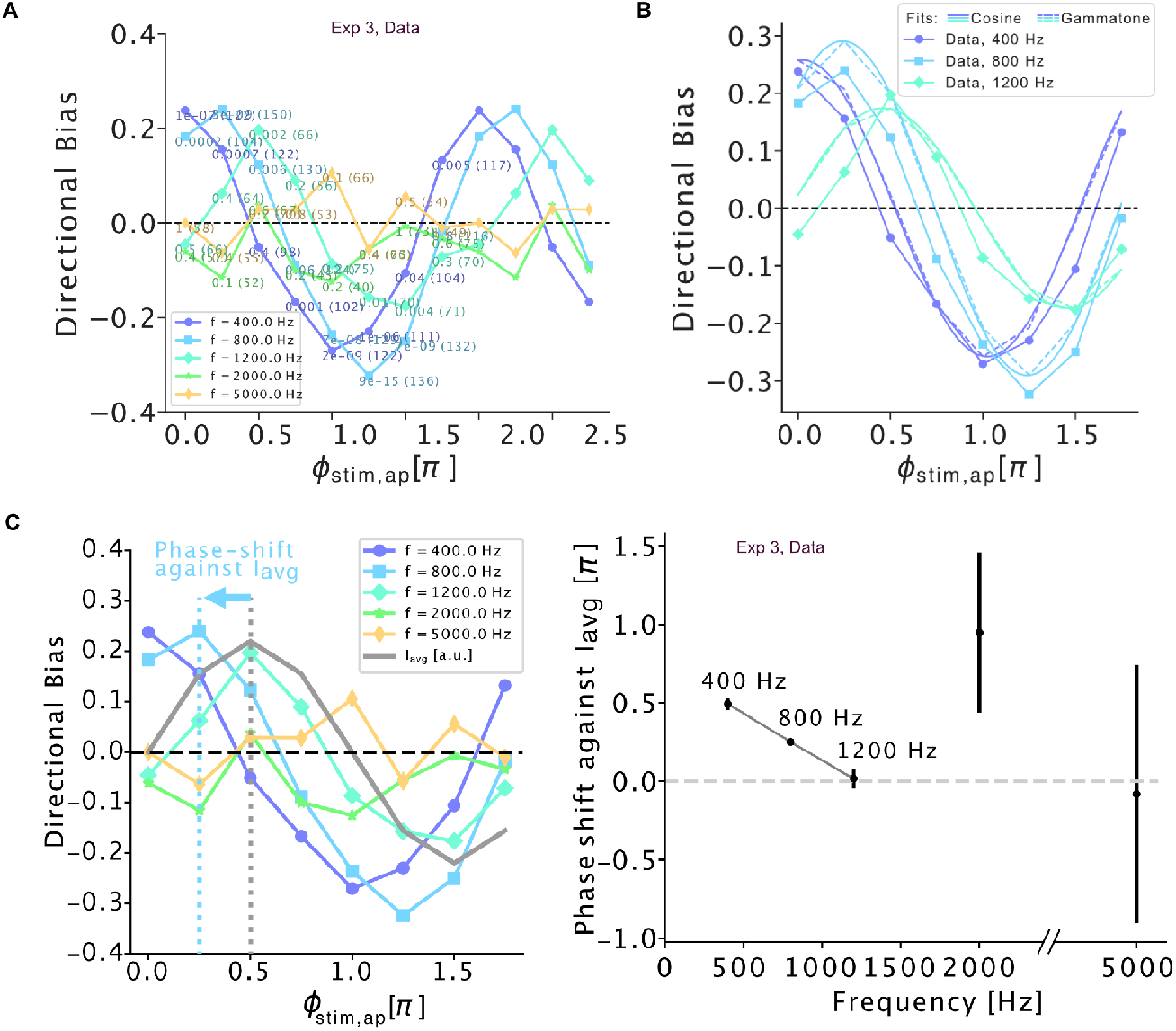
Directional bias. (A) Experimental data of experiment 3, same as in Figure 4C. Here, the significance of a non-zero directional bias (two-sided binomial test) is indicated for each data point, as well as the number of startles elicited in each stimulus condition (in brackets). 85 startles were evoked on average and at least 40 startles in each stimulus condition. (B) With the fish parameters fitted in Figure 4D, the generalized model fits the experimental data well (using τ_*bio*_ = 0. 27 *ms* ± 0. 02*ms* and ϕ_*bio,ap*_ = 0. 19 π ± 0. 04π), both using the analytical integral solution derived for cosine waveforms *D* ∝ *cos*(ϕ_*bio,vp*_ + ωτ_*bio*_ + ϕ_*stim,vp*_) and evaluating the integral numerically for the target gammatone waveforms used in the experiment. The amplitude was set to match the root mean square of the experimental data at each frequency. (C) The directional bias tuning to the relative phase Φ_stim,ap_ shifts linearly with frequency. Left: This phase shift is quantified by cross-correlating the tuning curves to the averaged acoustic intensity (grey). Right: Between 400 Hz and 1200 Hz, the phase shift against I_avg_ is linear in frequency, i.e., a time delay. The phase shifts are identified from the peaks in the cross-correlation. The error bars denote the standard deviation of the means, computed through bootstrapping: The tuning curves and the peaks of their cross-correlation with I_avg_ were recomputed 1000 times by resampling the original data with replacement.

**Figure S6.**
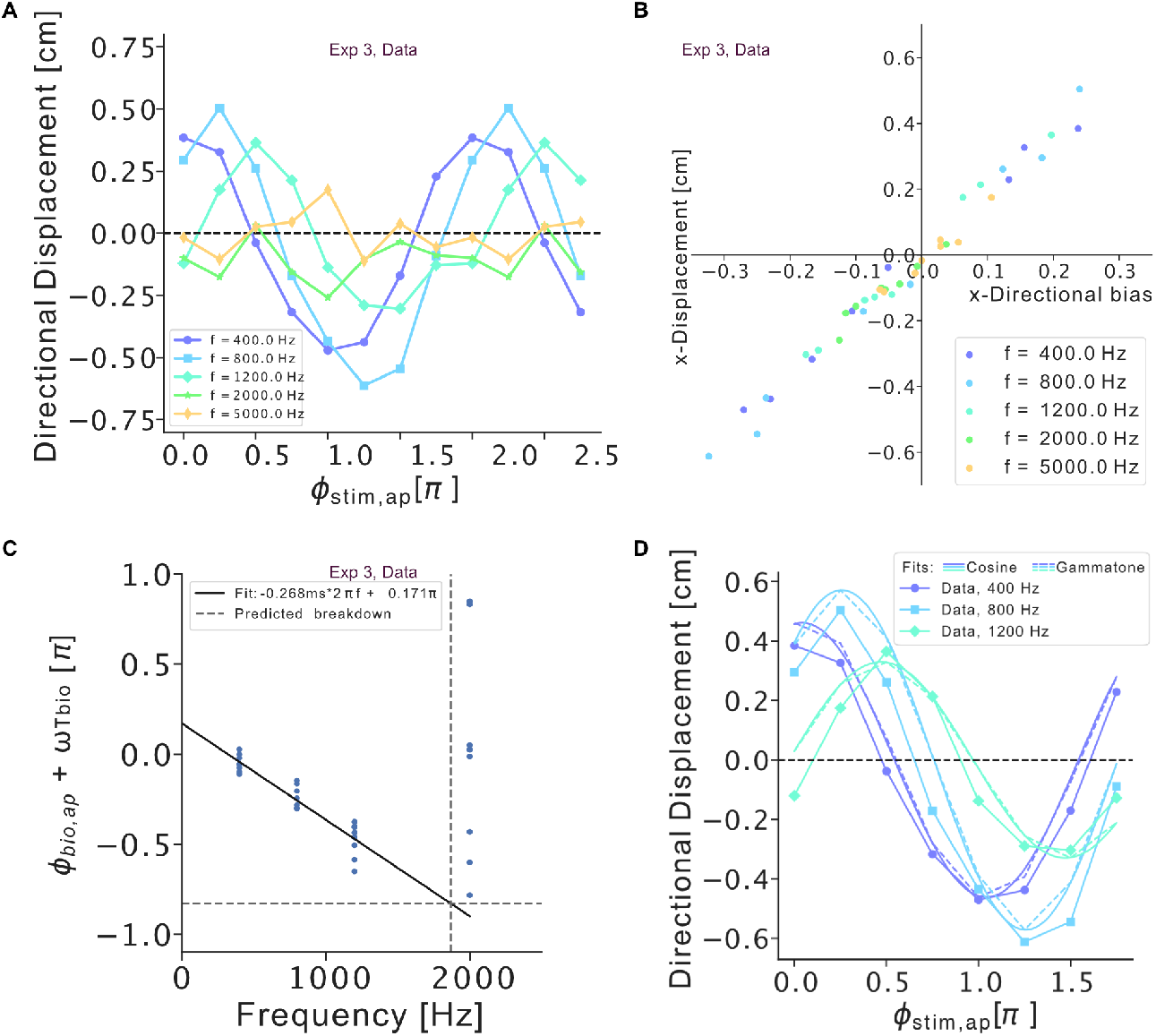
Directional displacement. (A) The tuning curves for directional startle displacement look very similar to the tuning curves for directional bias. (B) Directional bias and directional displacement are correlated. (C) Similar analysis to Figure 4D, repeated for startle displacement instead of directional bias. The obtained fish parameters, τ_*bio*_ = − 0. 27 *ms* ± 0. 02*ms* and ϕ_*bio,ap*_ = 0. 17π ± 0. 04π, are similar to the ones obtained from fitting directional bias (τ_*bio*_ = − 0. 27 *ms* ± 0. 02*ms* and ϕ_*bio,ap*_ = 0. 19 π ± 0. 04π). (D) Fit to data as in Figure S5B, here for directional displacement.

**Figure S7.**
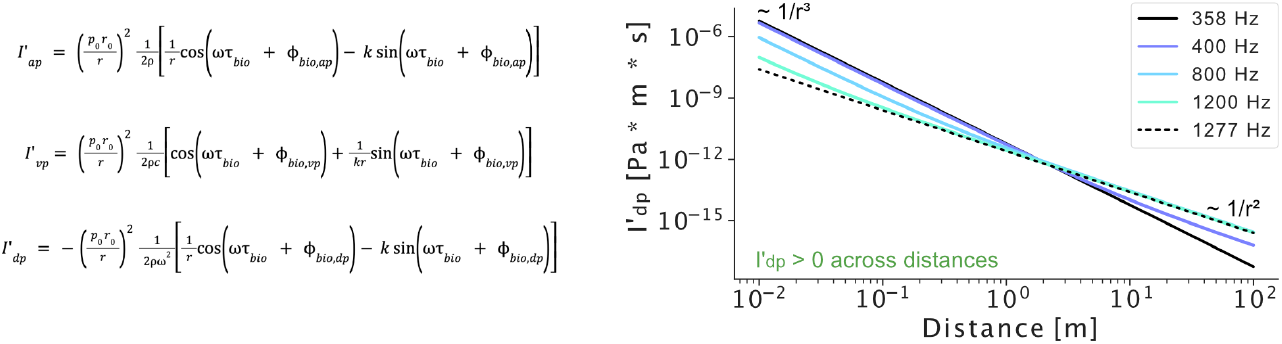
Model prediction across distances. The directional bias is computed analytically from 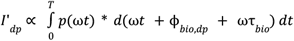 *dt*, using the sound monopole relationship between pressure and particle displacement for a sound of amplitude 147dB re. 1µPa at one body length (∼ 11 mm), and using ϕ_*bio,dp*_ = ϕ_*bio,ap*_ + π = 1. 19 π and τ_*bio*_ = − 0. 27 *ms* (Supplemental Information).

**Figure S8.**
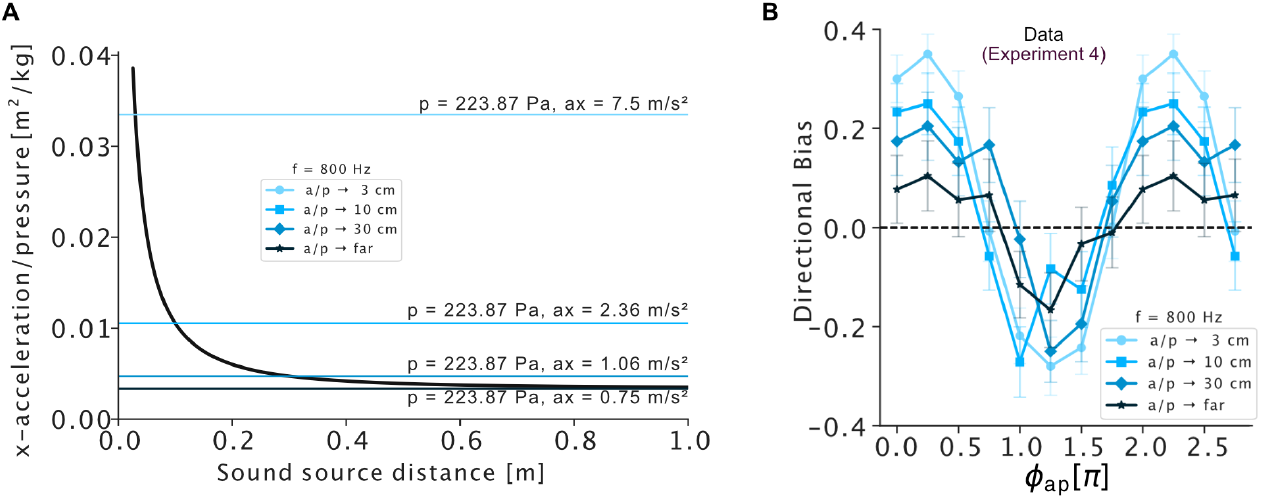
Directional bias tuning curves across amplitude ratios. (A) The amplitude ratio of particle acceleration and pressure is distance-dependent. (B) Tuning curves of directional bias with relative phase at f = 800 Hz for different amplitude ratios (a/p) of x-acceleration and pressure. Data from 3325 startles in 8885 playbacks across 133 fish. The phase-dependence of the directional bias is unaffected by the amplitude ratio. The pressure peak amplitude was fixed to 167 dB (re 1 µPa) and the particle acceleration amplitudes varied: 3 cm: 7.5 m/s^2^, 10 cm: 2.36 m/s^2^, 30 cm: 1.06 m/s^2^, Far: 0.75 m/s^2^. The error bars indicate the SEM.

